# The Indian Cancer Genome Atlas:A Multi-Omics Resource for Advancing Cancer Research

**DOI:** 10.1101/2025.03.25.645286

**Authors:** Suveera Dhup, Ankita Singh

## Abstract

Breast cancer genomics remains disproportionately shaped by cohorts of European and East Asian ancestry, leaving South Asian populations- nearly one-fifth of the world’s people-underrepresented despite a distinct clinical presentation: earlier age at onset, later stage at diagnosis, and poorer stage-specific survival relative to Western cohorts. Here we present an integrated multi-omic resource generated from the same 125-patient Indian breast cancer cohort, comprising whole-genome somatic variant calls, paired tumor–normal RNA-sequencing batches, and quantitative shotgun proteomics, together with clinical annotation spanning age range, menopausal status, tumor laterality, family history, receptor (ER/PR/HER2) status, and recurrence/progression outcome. Somatic profiling recovered canonical breast cancer drivers **(TP53, 36%; PIK3CA, 22%; GATA3, 8%; ARID1A, 5%)** alongside a set of recurrently altered genes dominated by exceptionally large genes (MUC4, TTN, OBSCN, USH2A), a recognized signature of length-driven passenger-mutation recurrence. Cross-referencing recurrent protein-level alterations against the OncoKB knowledge base identified PIK3CA/AKT1 pathway hotspot mutations, most frequently PIK3CA H1047R, in one-fifth of patients (25/125). Transcriptomic analysis cleanly separated tumors from normal tissue along the first principal component and supported a curated immune gene panel capable of stratifying tumors by immune-associated expression pattern. We provide this dataset as a resource for driver and actionability benchmarking, cross-platform (DNA-RNA-Protein) concordance analysis, tumor-immune stratification, and methodological work on length-normalized driver detection in an Indian-ancestry context that remains largely absent from reference cancer genomics datasets.

## Introduction

Breast cancer genomics has been shaped disproportionately by cohorts of European and East Asian ancestry. A comparative analysis of clinical and molecular data drawn from TCGA, the International Cancer Genome Consortium, and other major resources documented a significant paucity of South Asian ancestry data despite this group comprising up to one-fifth of the world’s population^1^. This is not a purely demographic gap: multi-omics profiling of Asian breast cancer cohorts has repeatedly surfaced population-specific patterns not captured by Western reference sets. A Malaysian cohort (MyBrCa, n=560) presented younger, at later stage, and with a higher proportion of HER2-positive tumors than TCGA^2^; a Korean cohort enriched for pre-menopausal patients showed a different molecular subtype mix, altered ESR1 expression, and - notably for the present dataset - GATA3 mutations enriched specifically in younger patients^3^; and an Indian transcriptomic study found subtype-specific mRNA/lncRNA signatures diverging from TCGA-derived patterns^4^. ICGA Foundation was established explicitly to address this gap, citing the characteristically younger age at diagnosis and later stage at presentation in Indian patients^5^.

These molecular differences track a distinct clinical picture on the ground. India recorded an estimated 200,000 new breast cancer cases in 2020, projected to rise to roughly 232,800 by 2025^6^. Indian series consistently report a median age at diagnosis around 47–49 years-roughly a decade younger than typical Western cohorts - with the large majority of cases clustering between 31 and 60 years of age^6^, and most hospital-registry cases present with locoregional or metastatic disease rather than localized tumors; population-based registry data similarly show a majority of Indian patients diagnosed at the locoregional stage, in contrast to the localized-stage predominance seen in the United States, with correspondingly lower stage-specific survival^7^.

Filling this gap requires more than a single-platform catalogue of variants. Somatic mutation calls alone cannot distinguish biologically meaningful driver events from passenger mutations inflated by gene length; transcriptomic data alone cannot confirm that expression differences are reflected at the functional (protein) level; and neither can establish whether a cohort’s tumor-immune landscape resembles or diverges from other populations. This dataset was therefore generated as an integrated resource: whole-genome somatic variant calls, paired tumor–normal RNA-sequencing, and quantitative proteomics from the same 125-patient Indian breast cancer cohort, together with clinical annotation suitable for stratified reuse. This Article accompanies the deposited dataset as a Data Descriptor-style resource paper: it does not report novel biological findings, but instead establishes, through technical validation against known breast cancer biology, that the resource is fit for reuse in hypothesis-driven analyses of an Indian breast cancer population.

## Results

### Cohort composition and clinical annotation

Whole-genome sequencing, RNA-sequencing, and proteomics were generated from the same 125- patient Indian breast cancer cohort. Clinical annotation was collected for each patient, including age range, smoking and alcohol history, tumor laterality, family history of cancer, menopausal status, estrogen receptor (ER), progesterone receptor (PR) and human epidermal growth factor receptor 2 (HER2) status, and recurrence/progression status. Patients were classified into four receptor-defined molecular subgroups: HER2-negative (n=57, 43.8%), triple-negative breast cancer (TNBC; n=29, 22.3%), HER2-positive (n=25, 19.2%), and triple-positive breast cancer (TPBC; n=19, 14.6%) (Fig. 1).

**Figure 1.**
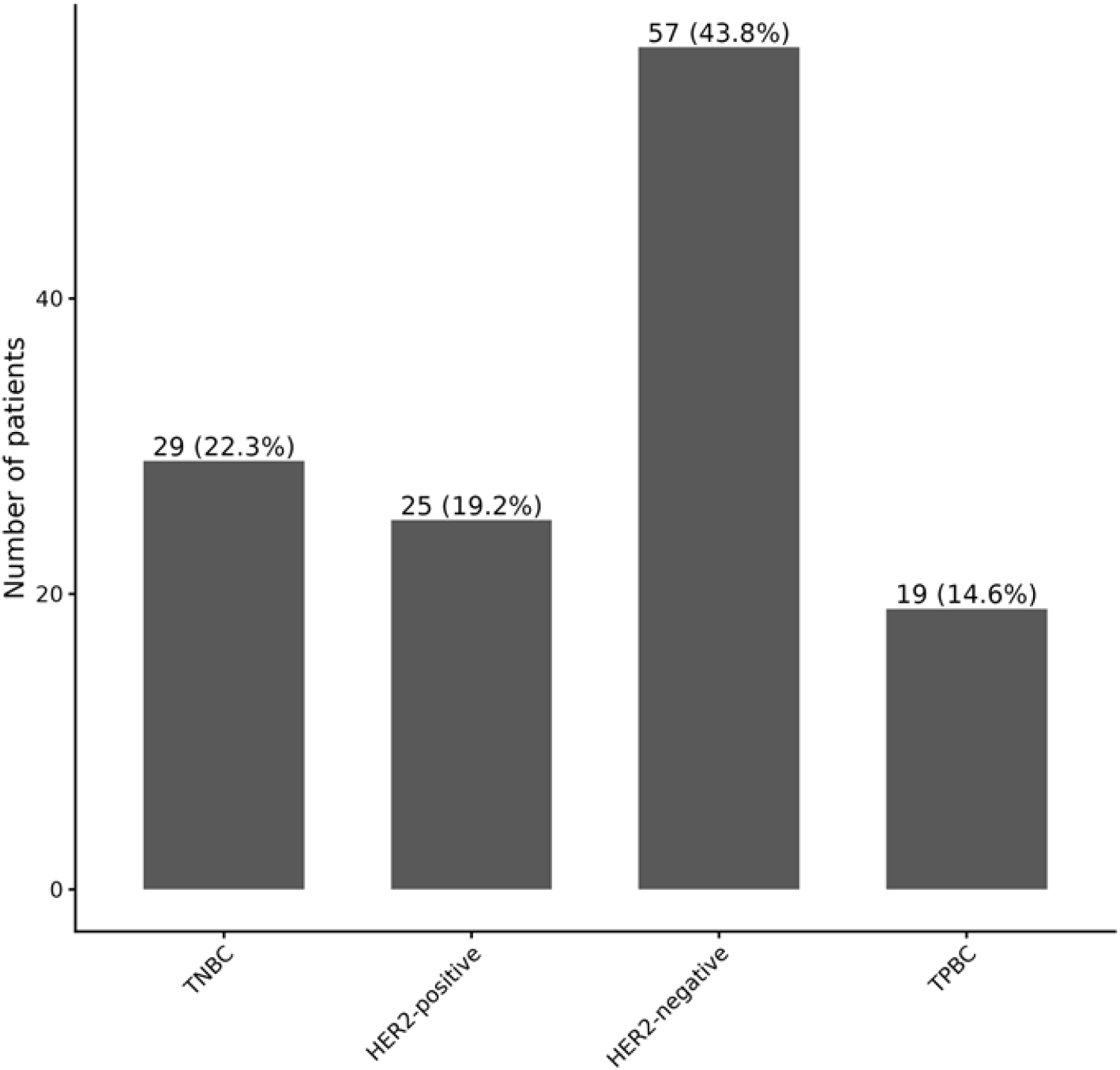
Distribution of receptor-defined molecular subtypes across the 125-patient cohort. Bar height indicates the number of patients per subtype; labels show patient count and percentage of the classified cohort. HER2-negative tumors were the largest subgroup, followed by TNBC, HER2-positive, and TPBC.

### Somatic mutational landscape

Individual sample-level MAF files generated from the whole-genome somatic variant-calling workflow were merged into a cohort-level variant dataset and characterized using maftools^9^. Missense mutations were the dominant functional class, single-nucleotide variants were the dominant alteration type, and C>T transitions were the most frequent single-nucleotide substitution class (n=2,437 of the combined substitution counts), consistent with the transition bias typically seen in solid tumor genomes (Fig. 2). The cohort had a median of 32 somatic variants per sample (range up to 268 in the plotted samples).

**Figure 2.**
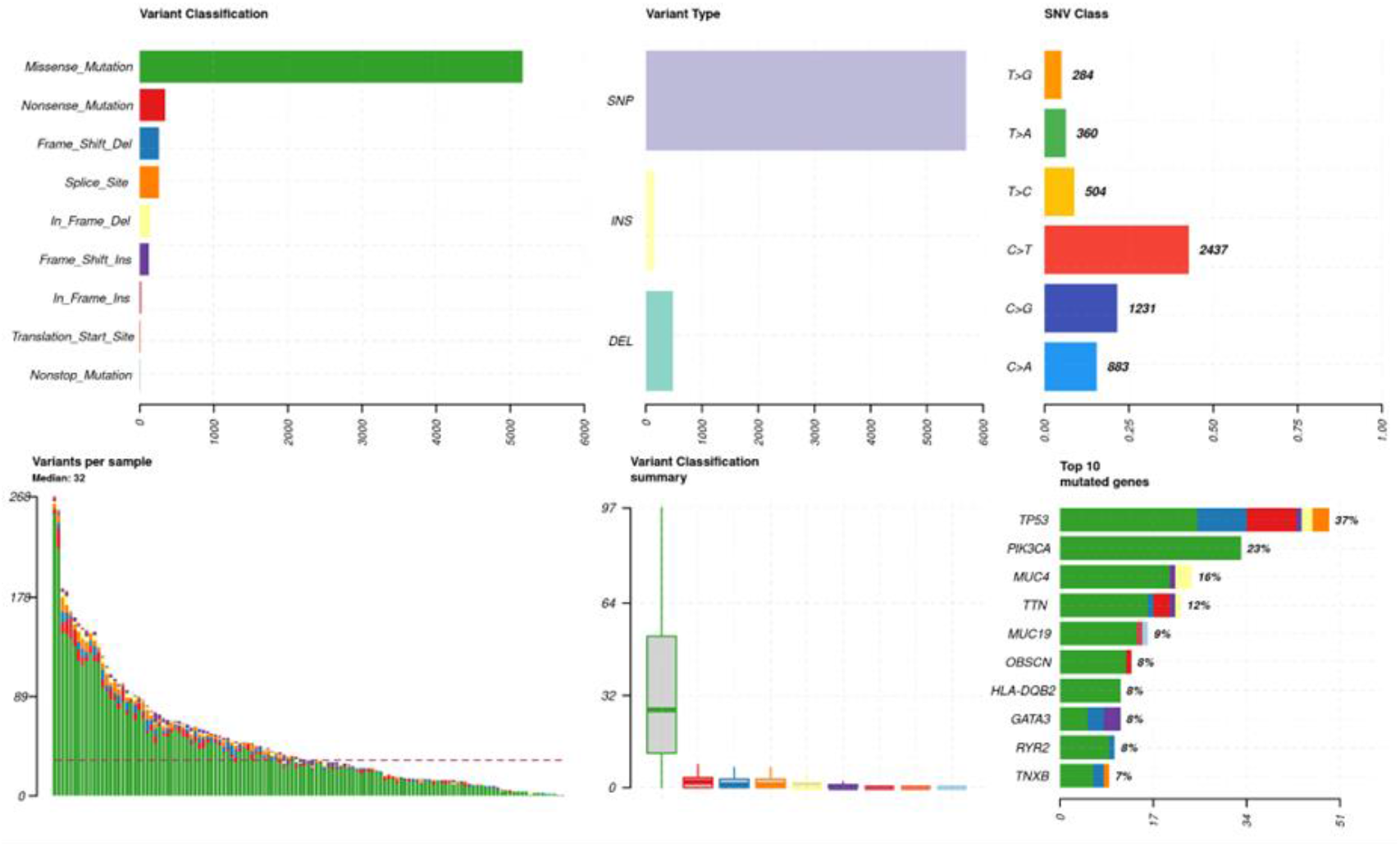
Cohort-level summary of somatic variant characteristics. (A) Variant classification by predicted functional consequence. (B) Variant type (SNP, insertion, deletion). (C) Singlenucleotide substitution class, with complementary substitutions combined by strand equivalence. (D) Variants detected per sample. (E) Per-sample variant counts by classification. (F) The ten most frequently mutated genes in the cohort, with the percentage of samples harboring an alteration in each gene.

The top mutated gene was TP53 (36% of samples), followed by PIK3CA (22%), MUC4 (16%), TTN (12%), MUC19 (9%), and a set of genes each altered in 5–8% of samples: GATA3, HLADQB2, OBSCN, RYR2, TNXB, AHNAK2, ARID1A, HMCN2, USH2A, CSMD1, FCGBP, HRNR, LRP2, PKHD1L1, and RYR1 (Fig. 3). TP53 and PIK3CA alterations are canonical breast cancer drivers and their prevalence in this cohort is broadly consistent with global breast cancer series. However, we note explicitly that many of the remaining recurrently altered genes in this list-MUC4, TTN, OBSCN, MUC19, USH2A, RYR2, CSMD1, HMCN2, LRP2, AHNAK2, TNXB, FCGBP, HRNR, RYR1, and PKHD1L1-are among the largest genes in the human genome (coding sequence lengths ranging from tens to over a hundred kilobases). Recurrent mutation of exceptionally large genes in exome- and genome-scale cancer cohorts is a well-documented artifact of gene length driving passenger-mutation accumulation rather than positive selection^14,15,16^, and their presence in this cohort’s raw top-20 list should not, by itself, be interpreted as evidence of oncogenic driver activity.

**Figure 3.**
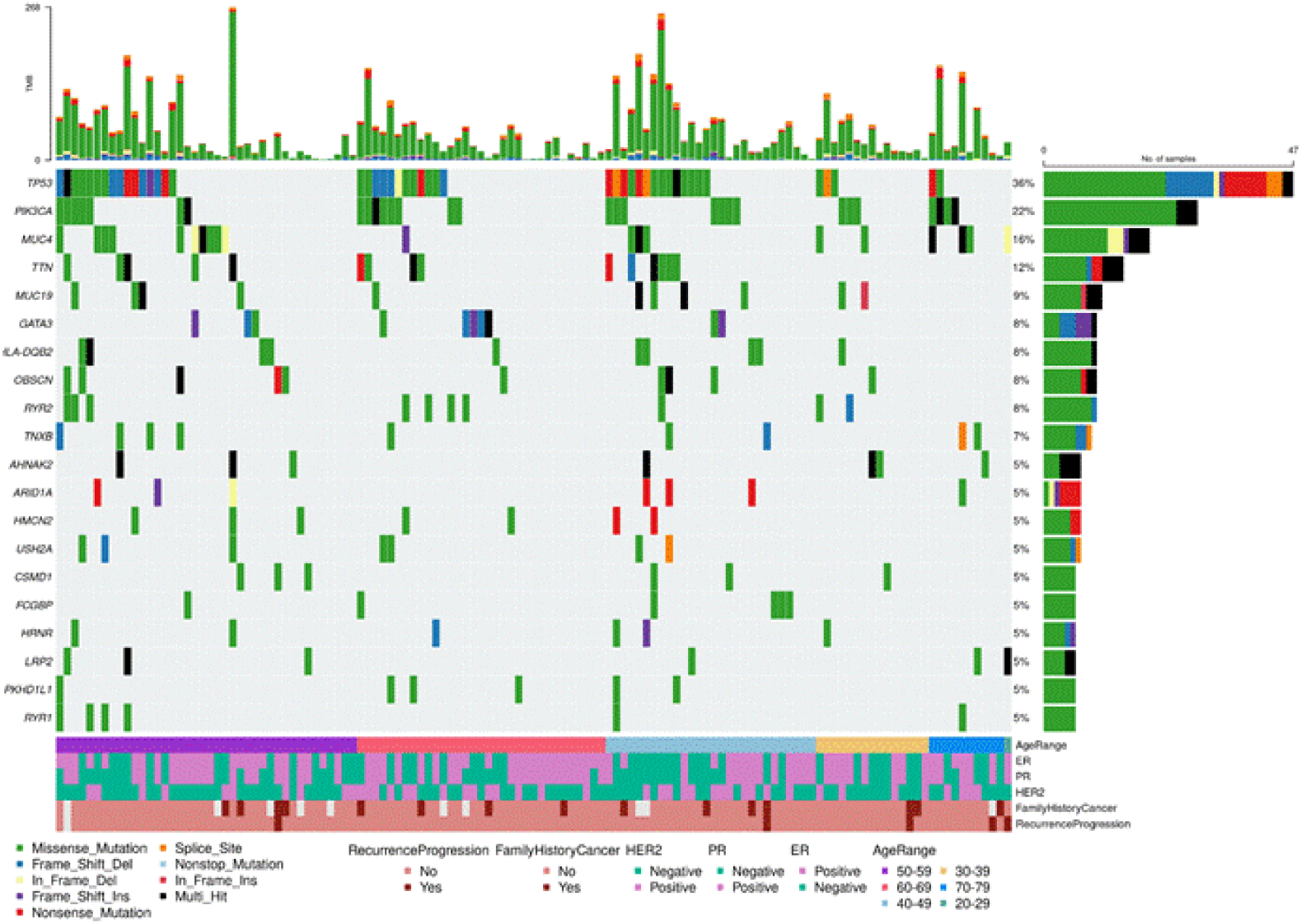
The somatic mutation landscape of the 20 most frequently mutated genes across the tumor cohort (oncoplot). Rows represent genes ranked by mutation frequency; columns represent individual tumor samples. Cell color denotes the variant classification. The top bar plot shows tumor mutational burden per sample; the right-hand bar plot shows mutation frequency per gene. Clinical annotation tracks (age range, ER, PR, HER2, family history of cancer, and recurrence/progression status) are shown below the mutation matrix.

Protein-level mapping of amino-acid substitutions (HGVSp_Short annotation) confirmed clustering of PIK3CA mutations at canonical hotspot codons (using reference transcript NM_006218) and of TP53 mutations across the DNA-binding domain (Supplementary Fig. S1– S2). Cross-referencing recurrent protein-level alterations against the OncoKB precision-oncology knowledge base^8^ identified eight recurrent gene–protein alterations with documented oncogenic and/or therapeutic annotation (Fig. 4). PIK3CA H1047R was the most common actionable alteration (13/125 patients, 10.4%), followed by PIK3CA E542K, E545K, and H1047L (3 patients each), PIK3CA C420R, H1047Y, and N345K (1 patient each), and a single AKT1 E17K alteration. In aggregate, 25 of 125 patients (20.0%) carried at least one OncoKB-annotated PIK3CA/AKT1 pathway hotspot mutation of potential relevance to PI3K-pathway-targeted therapy (e.g., alpelisib) under current companion-diagnostic criteria. This figure should not be extrapolated from Western cohorts given known population differences in PIK3CA mutation prevalence and receptor-subtype distribution, underscoring the value of population-specific actionability benchmarking.

**Figure 4.**
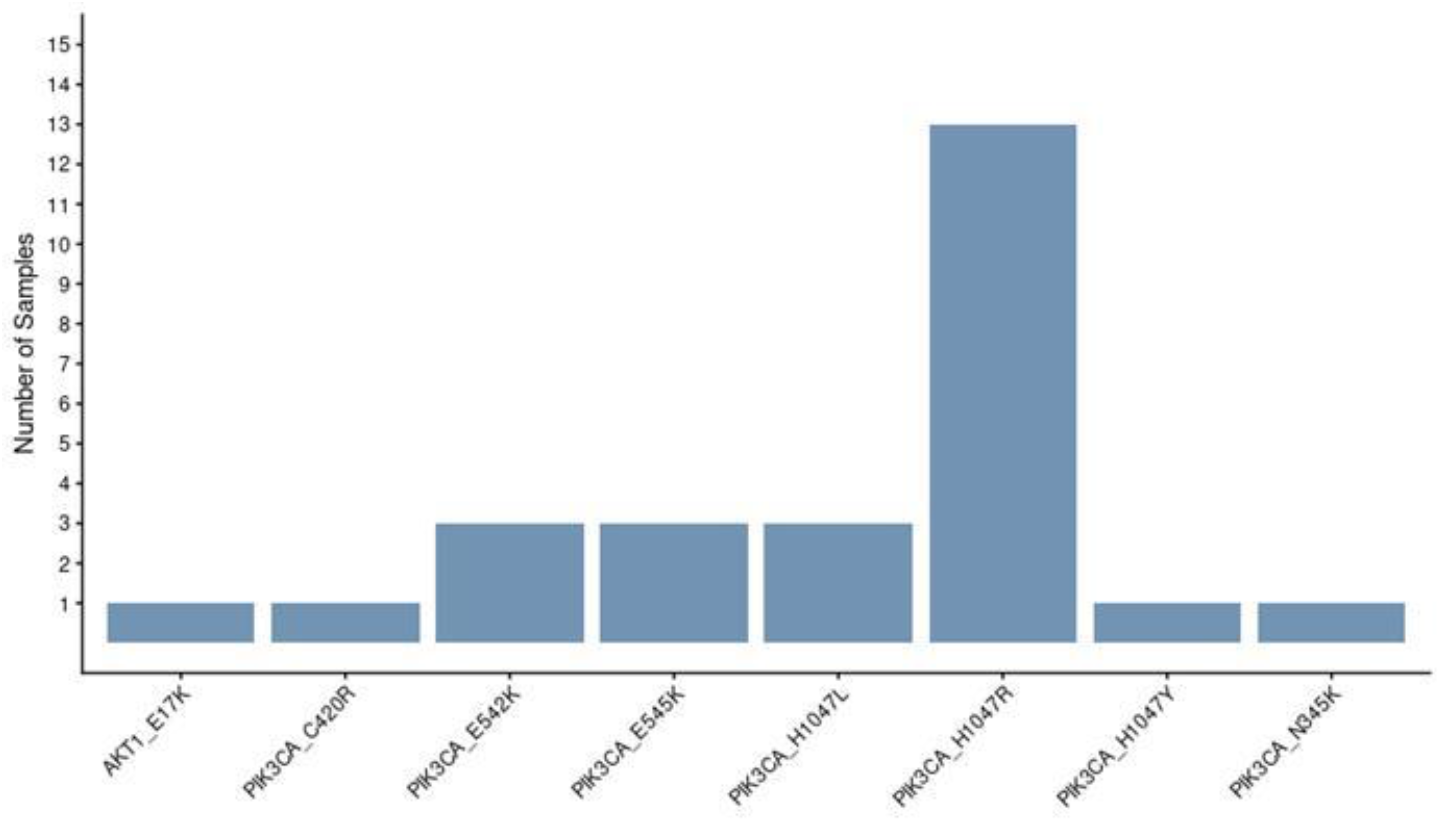
OncoKB-annotated potentially actionable somatic alterations. The x-axis denotes individual gene–protein alterations (gene symbol and amino-acid change); the y-axis denotes the number of patients harboring each alteration.

### Transcriptomic landscape

RNA-sequencing data were processed for 125 sample pairs. Transcript-level abundance was quantified with Salmon^10^ and summarized to gene level with tximport^11^ using a GENCODE hg38 transcript-to-gene annotation^13^. Differential expression between tumor and matched normal tissue was modeled with DESeq2^12^ using a paired design (∼Patient + Condition) to account for interpatient variability, with genes considered differentially expressed at a Benjamini–Hochberg adjusted P<0.05 and an absolute log2 fold-change >2. The125-sample batch showed a consistent overall pattern on MA-plot, volcano-plot, and PCA visualization (Supplementary Fig. S3–S5).

A curated immune gene panel spanning T-cell (CD3D/E/G, TRBC1/2), CD8 T-cell (CD8A, CD8B), cytotoxic/NK (NKG7, GNLY, GZMB, GZMH, PRF1, KLRD1, FCGR3A), B-cell/plasma-cell (CD79A, MS4A1, CD37, CD74, HLA-DRA, MZB1, JCHAIN, SDC1), monocyte/macrophage (LYZ, CD68, CSF1R, CTSS, FCER1G, TYROBP), dendritic-cell (CD1C, FCER1A, CLEC10A), neutrophil (S100A8, S100A9, FCGR3B, CSF3R), and immune-checkpoint (PDCD1, CD274, CTLA4, LAG3, TIGIT, HAVCR2, IDO1) markers was used to examine transcriptional immune signatures. Row-wise Z-score standardization and unsupervised hierarchical clustering of the full 125-sample panel (Fig. 5) resolved samples into immune-marker expression clusters, providing a basis for immune-hot/immune-cold stratification that can be tested against clinical annotation and against reports of elevated immune activity in other Asian breast cancer cohorts^2,3^.

**Figure 5.**
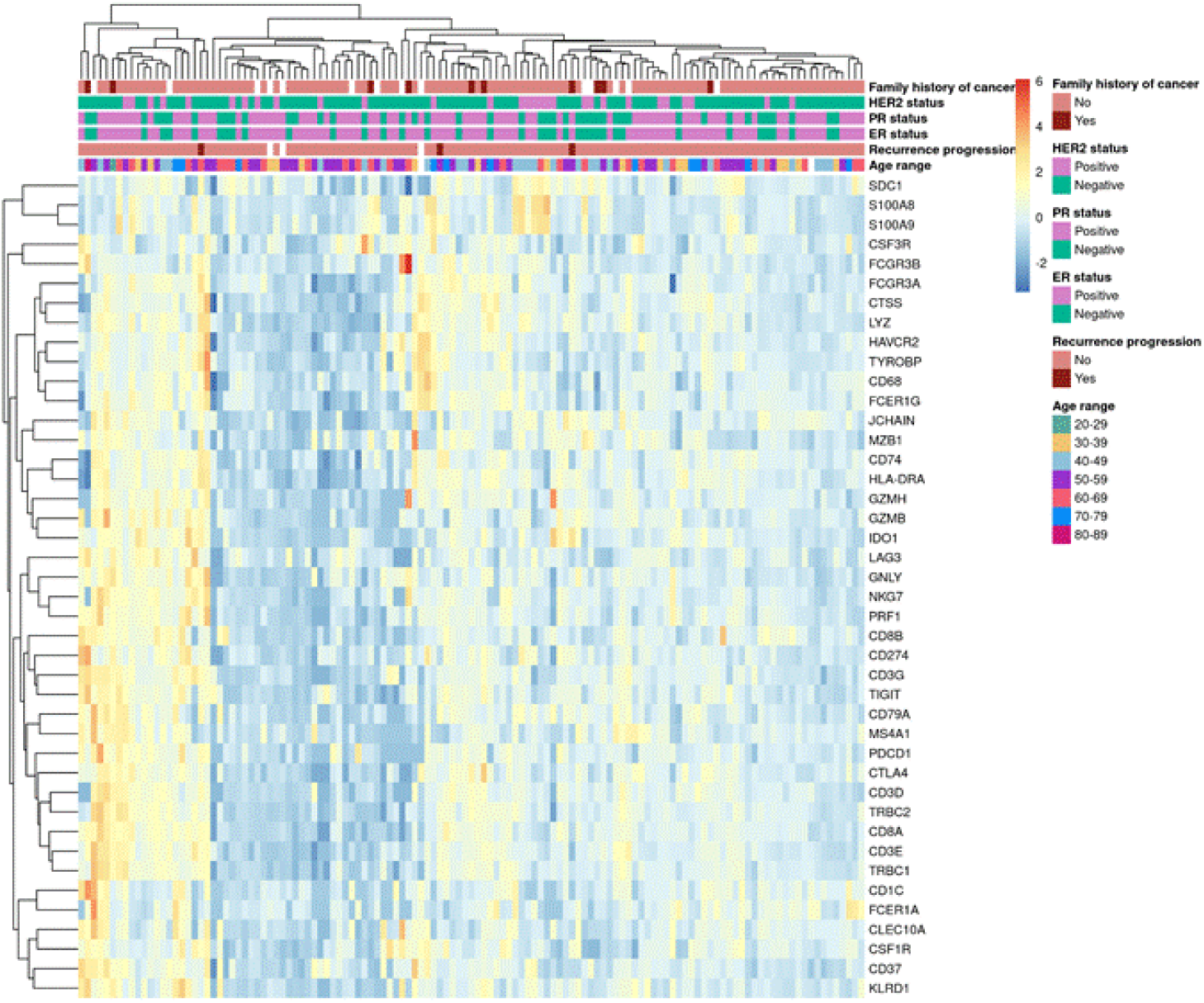
Immune-associated gene expression patterns across the 125-sample tumor RNA-seq cohort. Rows represent immune-associated genes; columns represent tumor samples, standardized by row-wise Z-score and hierarchically clustered. Column annotations indicate age range, ER, PR, HER2, family history of cancer, and recurrence/progression status.

The 20 genes most frequently mutated at the genomic level (Fig. 3) were cross-referenced against the independent RNA-seq dataset to examine their transcriptional profiles (Supplementary Fig. S7–S8, 48- and 125-sample subsets). This cross-platform view provides a direct, cohort-matched route for reusers to test whether recurrently mutated long genes also show coordinated expression change -a useful check when distinguishing candidate drivers from length-driven passenger events.

### Proteomic landscape

Quality-control-filtered, log2-normalized protein abundance profiles were generated for the tumor cohort and matched to clinical metadata and receptor-defined subgroup (TNBC, TPBC, HER2- positive, HER2-negative). Principal component analysis of the standardized abundance matrix showed that PC1 and PC2 (18.2% and 8.2% of variance, respectively) did not cleanly separate samples by receptor-defined subtype (Fig. 6); most samples formed a single dense cluster irrespective of subtype label, with a small number of outlier samples separating along PC1. Pairwise Pearson correlation and unsupervised average-linkage (UPGMA) hierarchical clustering of the same abundance matrix (Supplementary Fig. S9–S10) similarly grouped samples primarily by global proteomic similarity rather than by receptor subtype.

**Figure 6.**
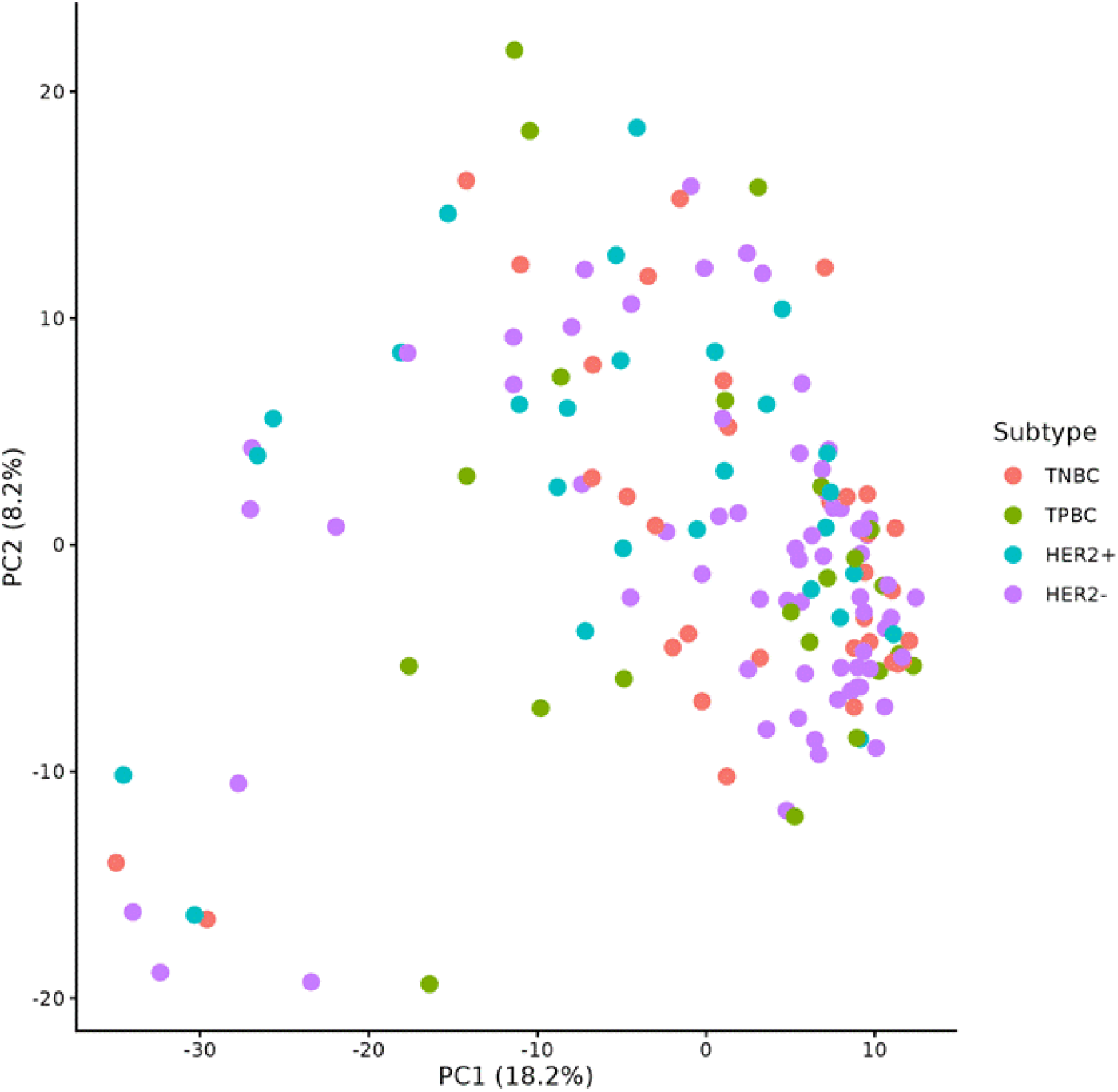
Principal component analysis of tumor proteomic profiles. Each point represents an individual tumor sample colored by receptor-defined subtype (TNBC, TPBC, HER2-positive, HER2-negative). PC1 and PC2 percentages indicate the proportion of total variance explained.

To characterize the dominant axes of proteomic variation independent of a specific classification scheme, the 250 proteins with the highest variance across QC-filtered samples were selected and visualized as a Z-score-standardized heatmap ordered by receptor-defined subtype (Fig. 7). This view revealed blocks of co-varying proteins associated with particular subtype groupings alongside substantial within-subtype heterogeneity, consistent with the incomplete subtype separation observed on global PCA.

**Figure 7.**
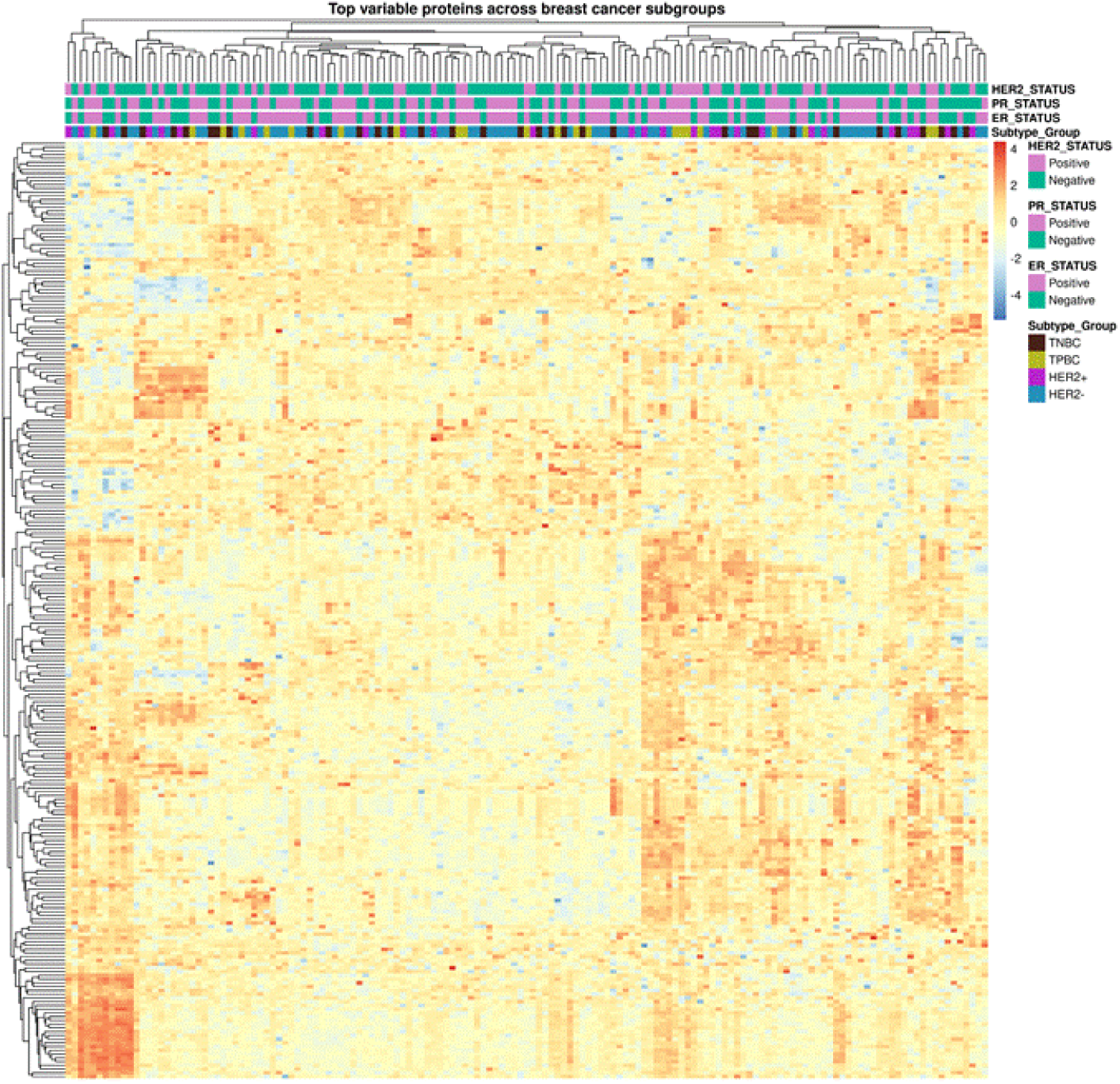
Heatmap of the 250 most variable tumor proteins. Columns represent tumor samples ordered by receptor-defined subtype (TNBC, TPBC, HER2-positive, HER2-negative); rows represent proteins clustered by abundance pattern. Color represents row-wise Z-score-transformed relative abundance. ER, PR, and HER2 status are shown as sample annotations.

To directly test whether transcriptional changes are reflected at the functional protein level, RNA-seq and proteomic measurements from the same tumors were cross-referenced at the gene level. A gene was called concordant if it reached significance independently in the proteomic comparison (P<0.05), in RNA-seq, and showed the same direction of change across all three measurements. This identified 481 concordantly altered genes (195 upregulated, 286 downregulated in tumor relative to normal tissue), which we provide as a supplementary table for reuse.

The concordant gene set resolved into distinct, biologically interpretable blocks. The largest and most consistent axis was a loss of adipocyte and lipid-metabolism markers in tumor tissue (PLIN4, FABP4, CD36, AOC3, ADH1B, GPD1, CES1, AKR1C2, CRYAB, HSPB6, CAV1, SOD3), with log2 fold-changes reaching −3.6 at the protein level and below-8 at the RNA level -consistent with tumor tissue displacing the adipocyte-rich normal breast parenchyma used as the comparator. A second, opposing block comprised concordantly upregulated luminal/epithelial keratins (KRT7, KRT8, KRT18, KRT19) alongside extracellular matrix and stromal-remodeling genes (POSTN, FN1, VCAN, AEBP1, COL1A1, COL4A1, COL4A2, COL5A1, COL15A1), consistent with disrupted ductal architecture and a desmoplastic stromal reaction. A third block comprised concordantly upregulated spliceosome and RNA-binding proteins (SRSF2, SRSF3, SRSF6, HNRNPA1, HNRNPA2B1, HNRNPC, HNRNPD, HNRNPH1, HNRNPK, HNRNPL, PTBP1, NONO, SFPQ, DHX9, DHX15), consistent with the increased RNA-processing demands of proliferating tumor cells.

In parallel, comparison of protein- and RNA-level changes surfaced a distinct discordance pattern within a single, well-defined gene family: of 68 detected ribosomal protein genes (RPL/RPS family), all 68 were significantly increased at the protein level (67/68 with positive fold-change), but only 31/68 reached significance at the RNA level in the 250-sample batch, with several trending flat or slightly negative (e.g., RPL37A, RPS25, RPS6, RPL24, RPL31). This proteinup/RNA-flat pattern across an entire gene family is consistent with increased ribosome biogenesis or translational output in tumor cells that is not fully explained by transcriptional upregulation and illustrates the kind of post-transcriptional regulation that single-platform (RNA-only) analyses of this cohort would miss.

### Portal for data sharing

To support data sharing and enable global collaborations, ICGA has launched a dedicated **multiomics data portal** (https://www.icga.in/icga-portal/, https://icga.net.in) built on the cBioPortal framework. The ICGA Data Portal integrates clinical data with molecular information, aiming to serve as a valuable resource for precision oncology. This user-friendly platform offers seamless visualization, integration, and analysis of genomic, transcriptomic, proteomic, and clinical datasets. By facilitating open yet controlled access, the portal empowers researchers, clinicians, and policymakers to explore and utilize the data for biomarker discovery, treatment optimization, and translational research. Physicians may leverage this platform to tailor treatments based on genetic vulnerabilities by examining clinical outcomes and treatment regimens of patients with similar alterations. Additionally, researchers and clinicians can contribute genomic data, expanding the dataset for broader comparisons and insights. As a comprehensive cancer “knowledge bank,” the ICGA plays a crucial role in advancing precision oncology strategies. The portal will continue to grow more with data, clinical information on breast cancer, and many more cancer types.

## Discussion

This resource recapitulates expected breast cancer biology while surfacing features specific to an understudied population. The recovery of TP53 and PIK3CA as the two most recurrently altered genes, at frequencies broadly consistent with global breast cancer series, together with clean tumor-versus-normal separation on transcriptomic PCA, indicates that the whole-genome and RNA-sequencing components of this dataset behave as expected against known breast cancer biology and are fit for downstream reuse. At the same time, the dominance of exceptionally large genes among the next tier of recurrently altered genes is a reminder that raw mutation-frequency rankings, taken at face value, will overstate the driver landscape in any newly sequenced cohort - a risk that is arguably higher, not lower, in an Indian-ancestry cohort where established driver gene lists and length-normalization baselines have been developed almost entirely on Western and East Asian data. We provide the underlying per-gene mutation and coding-sequence-length data needed for others to apply, compare, or improve upon length-normalized driver-detection methods (e.g., MutSigCV^14^, dNdScv^15^, OncodriveFML^16^) specifically in this population.

The OncoKB-annotated actionability analysis illustrates why population-specific benchmarking matters in practice, not only in principle: one-fifth of this cohort carried a PIK3CA/AKT1 pathway hotspot alteration of potential relevance to PI3K-pathway-targeted therapy. Whether this proportion, and the specific hotspot distribution (dominated by PIK3CA H1047R), matches or diverges from Western and other Asian cohorts is precisely the kind of question that requires a directly comparable, population-matched dataset rather than extrapolation, and this resource is intended to support that comparison.

The proteomic results add a further, non-trivial observation: receptor-defined molecular subtypes that are well separated by design (ER/PR/HER2 immunohistochemistry) and reasonably well reflected in transcriptomic data did not resolve into distinct clusters on global proteomic PCA. We present this as an open empirical finding rather than a negative result to set aside - it may reflect genuine biological convergence of proteomic states across receptor subtypes, or a small number of outlier samples dominating the leading principal components, and we encourage users to investigate it directly using the deposited abundance matrices.

Taken together, we anticipate four primary reuse cases for this resource. First, driver and actionability benchmarking: recurrent alterations in PIK3CA, TP53, GATA3, and ARID1A can be cross-referenced against precision-oncology knowledge bases using the OncoKB-annotated actionability table provided with this resource. Second, cross-platform concordance benchmarking: because DNA, RNA, and protein-level measurements were generated from the same tumors, independent groups can test their own concordance pipelines. Third, immune**-** Microenvironment stratification: the curated checkpoint- and lineage-focused immune gene panel enables immune-hot/immune-cold subtyping in a population where such data remain sparse and allows direct testing of whether the more immune-active microenvironments reported in some younger Asian breast cancer cohorts^2,3^ generalize to an Indian cohort. Fourth, methodological benchmarking for long-gene passenger-mutation correction: the raw top 20 mutated-gene list, dominated by some of the largest genes in the human genome, together with the per-gene mutation and coding-sequence-length data provided, offers a population-specific test case for length-normalized driver-detection methods.

## Methods

### Cohort, clinical annotation, and ethics

All patients who provided informed consent for the study were between 18 and 75 years of age, treatment-naïve, and underwent whole-genome sequencing at a depth of 70X (±10), along with total RNA sequencing and proteomic analysis. For each patient, breast-adjacent non-tumor tissue was included as a control. The study was approved by the IECs of all contributing hospitals.

Sample quality was ensured through rigorous quality control measures at each step. Only samples with a minimum tumor content of 50%, as determined by a pathologist, were included in the study. Tumor purity was further assessed using bioinformatic tools applied to sequencing data. Clinical variables collected included cancer type, sex, age range, smoking and alcohol history, tumor laterality, family History of cancer, menopausal status, tobacco exposure, ER, PR, and HER2 status, and recurrence/progression status. The patients were followed for at least 3 years. Patient identifiers in the clinical metadata were matched to tumor sample barcodes using the numeric component of the corresponding identifiers.

### Whole-genome sequencing and somatic variant analysis

Whole-genome sequencing was performed *on Illumina Novaseq X Plus using manufacture’s protocol. Each sample was sequenced using 2×150bp chemistry, having a depth of 60-70X*, and somatic variants were called. Individual sample-level MAF files were read into R using data. Table and concatenated, retaining all available columns across samples, to produce a cohort-level somatic variant dataset. The merged dataset was imported into maftools^9^ for variant summarization, oncoplot generation, and lollipop-plot visualization. For the oncoplot, the 20 most frequently mutated genes were displayed with clinical annotation tracks (cancer type, sex, age range, smoking/alcohol history, tumor laterality, family history of cancer, menopausal status, tobacco exposure, ER, PR, HER2 status, and recurrence/progression status) shown above the mutation matrix. For protein-level (lollipop) visualization, amino-acid substitution annotations were obtained from the HGVSp_Short field and plotted against protein position; for PIK3CA, transcript reference NM_006218 was specified to ensure consistent protein-level mapping.

### Actionability annotation

Recurrent protein-level somatic alterations were cross-referenced against the OncoKB precision-oncology knowledge base^8^ to identify alterations with documented oncogenic and/or therapeutic annotation. Alteration-level patient counts were tabulated and visualized as a bar plot of gene– protein alteration versus number of patients harboring each alteration.

### RNA-sequencing and differential expression analysis

Transcript-level abundance estimates were generated with Salmon^10^ and imported into R using tximport^11^, summarizing transcript-level estimates to gene level using a transcript-to-gene mapping derived from the GENCODE hg38 annotation^13^. Differential expression between tumor and matched normal tissue was assessed using a paired design (∼Patient + Condition, with patient included as a blocking factor) under the negative-binomial framework implemented in DESeq2^12^. Multiple-testing correction was performed using the Benjamini–Hochberg procedure; genes were considered differentially expressed at an adjusted P<0.05 and an absolute log2 fold-change >2, with log2 fold-change >2 classified as tumor-upregulated and <−2 as tumor-downregulated. For exploratory visualization and PCA, variance-stabilizing transformation (DESeq2 vst, blind=FALSE) was applied to the filtered dataset, and the percentage of variance explained by the first two principal components was used to describe the contribution of each component to overall sample variation.

### Immune gene panel analysis

A predefined immune-associated gene panel (see Results) was examined using the variance-stabilized expression matrix. Gene identifiers were mapped to gene symbols using the corresponding GENCODE annotation. Expression values for each gene were standardized across tumor samples by row-wise Z-score, and hierarchical clustering was performed independently for genes and samples using the standardized matrix. Clinical annotations (age range, ER, PR, HER2 status, family history of cancer, and recurrence/progression status) were incorporated as column annotations.

### Cross-referencing WGS-derived genes with RNA-seq expression

The 20 genes most frequently mutated in the WGS oncoplot were selected and their expression examined in the independent RNA-seq dataset using the same variance-stabilization and Z-score standardization approach described above, with hierarchical clustering of genes and samples and the same clinical annotation tracks.

### Proteomics data processing and analysis

Protein abundance data were quality-control-filtered and log2-normalized. Samples were matched to clinical metadata and classified into receptor-defined subgroups based on ER, PR, and HER2 status. For visualization, the QC-filtered, log2-normalized abundance matrix was standardized independently for each protein across samples using row-wise Z-score transformation. Hierarchical clustering was performed independently for proteins and samples using the standardized matrix. Principal component analysis was performed on the same matrix, with samples colored by receptor-defined subgroup. Pairwise sample similarity was additionally assessed using Pearson correlation coefficients calculated from the QC-filtered, log2-normalized abundance matrix; a distance matrix was generated as 1−r, and samples were clustered using average-linkage (UPGMA) hierarchical clustering. For the top-variable-protein analysis, protein-wise variance was calculated across all QC-filtered samples and the 250 highest-variance proteins were selected and visualized as a row-wise Z-score-standardized heatmap ordered by receptor-defined subtype.

## References

1. The clinical and molecular landscape of breast cancer in women of African and South Asian ancestry. medRxiv 2024.05.15.24307435 (2024).

2. The molecular landscape of Asian breast cancers reveals clinically relevant population-specific differences. Nat. Commun. 12, 32 (2021).

3. The Indian Cancer Genome Atlas: A Multi-Omics Resource for Advancing Cancer Research. bioRxiv 2025.03.25.645286 (2025).

4. National Academy of Medical Sciences (India) Task Force Report on Breast Cancer in India. Ann. Natl. Acad. Med. Sci. (India) 61, 132–170 (2025).

5. Sathishkumar, K. et al. Breast cancer survival in India across 11 geographic areas under the National Cancer Registry Programme. Cancer 130, 2181–2192 (2024).

6. Chakravarty, D. et al. OncoKB: A Precision Oncology Knowledge Base. JCO Precis. Oncol. 1, 1–16 (2017).

7. Mayakonda, A., Lin, D. C., Assenov, Y., Plass, C. & Koeffler, H. P. Maftools: efficient and comprehensive analysis of somatic variants in cancer. Genome Res. 28, 1747–1756 (2018).

8. Patro, R., Duggal, G., Love, M. I., Irizarry, R. A. & Kingsford, C. Salmon provides fast and bias-aware quantification of transcript expression. Nat. Methods 14, 417–419 (2017).

9. Soneson, C., Love, M. I. & Robinson, M. D. Differential analyses for RNA-seq: transcript-level estimates improve gene-level inferences. F1000Research 4, 1521 (2015).

10. Love, M. I., Huber, W. & Anders, S. Moderated estimation of fold change and dispersion for RNA-seq data with DESeq2. Genome Biol. 15, 550 (2014).

11. Frankish, A. et al. GENCODE reference annotation for the human and mouse genomes. Nucleic Acids Res. 47, D766–D773 (2019).

12. Lawrence, M. S. et al. Mutational heterogeneity in cancer and the search for new cancer-associated genes. Nature 499, 214–218 (2013).

13. Martincorena, I. et al. Universal patterns of selection in cancer and somatic tissues. Cell 171, 1029–1041 (2017).

14. Mularoni, L., Sabarinathan, R., Deu-Pons, J., Gonzalez-Perez, A. & López-Bigas, N. OncodriveFML: a general framework to identify coding and non-coding regions with cancer driver mutations. Genome Biol. 17, 128 (2016).

